# Discovering subgenomes of octoploid strawberry with repetitive sequences

**DOI:** 10.1101/2020.11.04.330431

**Authors:** Adam Session, Daniel Rokhsar

## Abstract

Although its sequence was recently determined in a genomic *tour de force*,{Edger 2019} the ancestry of the cultivated octoploid strawberry *Fragaria* x *ananassa* remains controversial.{Liston 2020; Edger 2020} Polyploids that arise by hybridization generally have chromosome sets, or subgenomes, of distinct ancestry.{Stebbins 1947; Garsmeur 2014} The conventional method for partitioning a polyploid genome into its constituent subgenomes relies on establishing phylogenetic relationships between protein-coding genes of the polyploid and its extant diploid relatives,{Edger 2018-sub} but this approach has not led to a consensus for cultivated strawberry.{Liston 2020; Edger 2020} Here we resolve this controversy using a complementary strategy that focuses on the chromosomal distribution of transposable elements and depends only on the octoploid sequence itself.{Session 2016; Mitros 2020} Our method independently confirms the consensus that two of the four subgenomes derived from the diploid lineages of *F. vesca* and *F. iinumae*.{Tennessen 2014; Edger 2019} For the remaining two subgenomes, however, we find a statistically well-supported partitioning that differs from ref. {Edger 2019} and other work (reviewed in {Hardigan 2020}). We also provide evidence for a shared allohexaploid intermediate and suggest a neutral explanation for the “dominance” of the *F. vesca*-related subgenome.

---

Octoploid strawberries exhibit disomic inheritance (2n = 8x = 56).{Tennessen 2014; van Dijk 2014; Hardigan 2020} This suggests that the chromosome sets derived from different progenitors may persist as distinct subgenomes, albeit with a modest level (∼11%) of homeologous exchange.{Edger 2019; Edger 2020} Allopolyploid formation (i.e., hybridization followed by genome doubling) brings together chromosome sets with distinct evolutionary histories into the same nucleus.{Stebbins 1947; Garsmeur 2014} Since transposable elements expand and spread across nuclear genomes in irregularly timed bursts of activity,{Dangel 1995; Ma 2004} the chromosomal distribution and timing of transposon insertions provides a novel tracer of chromosome history in polyploids.{Session 2016; Mitros 2020} Importantly, this history is recorded in the genome of the polyploid itself, and can be recovered without reference to extant diploids. This is particularly important when the relevant diploid lineages are extinct.

**Figure 1a** shows that the octoploid strawberry chromosomes can be naturally partitioned into four distinct subgenomes, each of which is a complete chromosome set (*i*.*e*., one chromosome from each homeologous quartet) with a characteristic pattern of past repetitive activity. This partitioning is evident from 829 13 bp sequences (13-mers) that are (1) present in at least 100 copies in the octoploid, and so tag repetitive elements, and (2) are two-or-more-fold enriched on at least one chromosome from each homeologous quartet (**Suppl. Note 1; Ext. Data Fig 1; Suppl. Data File 1**). Importantly, our analysis is agnostic to prior hypotheses about subgenome identity.{Edger 2019; Liston 2020} **Fig. 1a** also includes the two available chromosome-scale diploid strawberry genomes, *F. vesca*{Edger 2018-ves} and *F. iinumae*.{Edger 2020} Although the 13-mers shown are defined only based on the octoploid genome, they also group each diploid set together.

**Figure 1.**
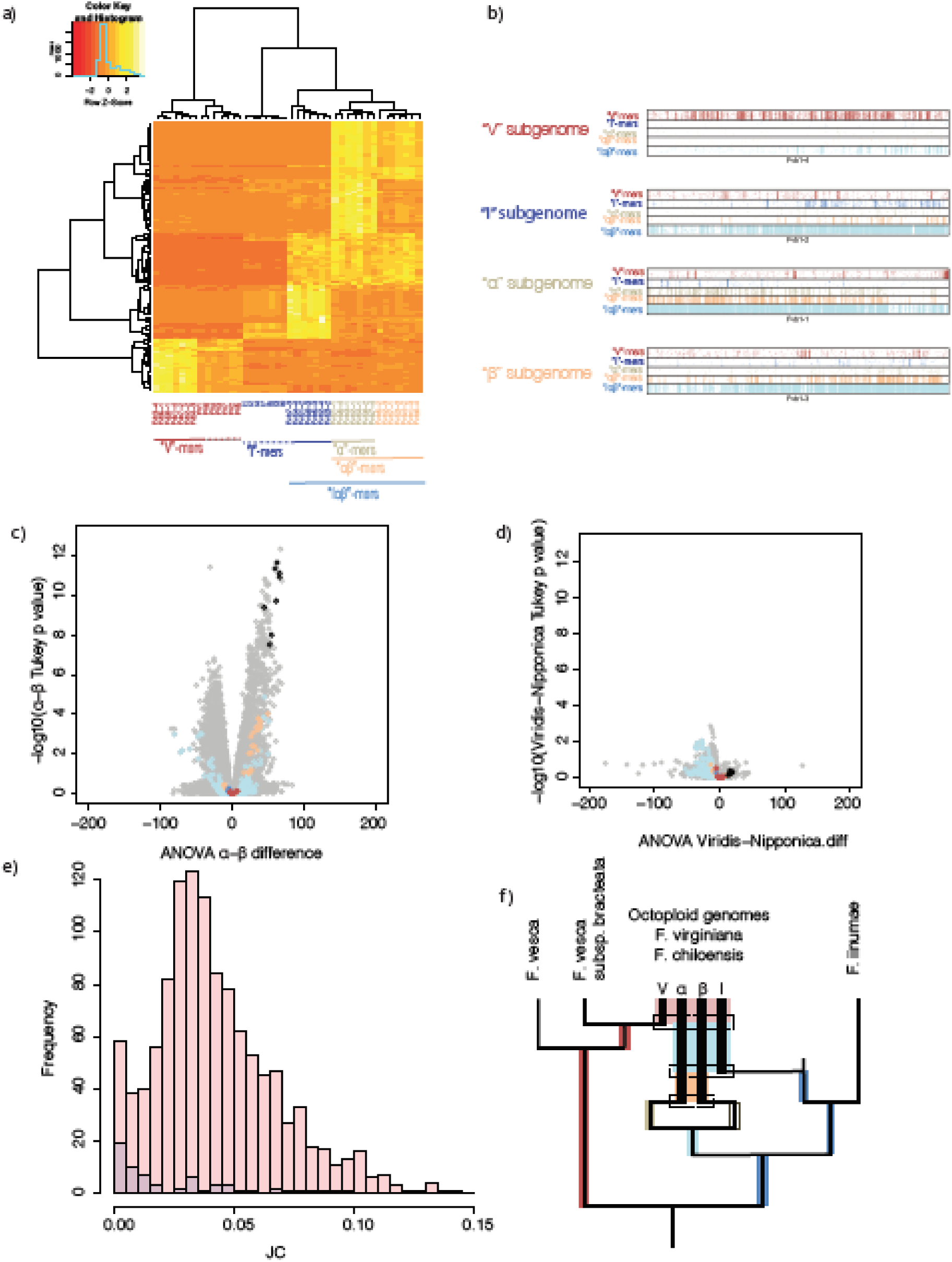
Repetitive sequences partition the octoploid genome into distinct subgenomes. **a)** 13-mers enriched in one or more chromosomes of each homeologous quartet consistently partition the octoploid genome into four distinct subgenomes: V, I, *α*, and β. Chromosomes are numbered according to Edger et al. [Edger 2019]. We chose 25 13mers of each type for visualization to allow the weaker I, *α*β, and *α* signals to be shown with the stronger V and I*α*β signals. Full dataset shown in **Ext. Data Fig. 1**. **b)** Distribution of four classes of 13-mer (V, I, *α, α*β, I*α*β) across a quartet of homeologous chromosomes of octoploid strawberry (chromosome 1 shown; others shown in **Ext. Data Fig. 2**). Short regions of homeologous exchange are evident. **c)** Volcano plot showing ANOVA *α*-β difference on x-axis, -log10(*α*-β Tukey p value) on the y. Black dots correspond to *α*-enriched 13mers, orange is *α*β, light blue is I*α*β, dark blue is I, and red is V. As expected the *α*-enriched 13mers are significantly higher in *α* than β (p<1e-7), and the I and V-specific 13mers are not informative for this comparison. **d)** Volcano plot showing ANOVA ‘Viridis’-’Nipponica’ difference on the x-axis, -log10(‘Viridis’-’Nipponica’ p-value) on the y. Colored dots are identical to c). There are no 13mers that differentiate these two chromosome sets. **e)** Histograms Jukes-Cantor distance between 5’ and 3’ LTRs for LTR-retrotransposons in large families with I*α*β-enriched 13-mers. Red: LTRs from I*α*β families on I*α*β chromosomes; blue: LTRs on V chromosomes. The peak at ∼0.035 (corresponding to ∼3 Mya, see **Suppl. Note 3**) implies coexistence of subgenomes at that time. Recent activity ∼1 My represent reactivations of this family induced by octoploid formation. This recent activity is found in a 3:1 ratio on I*α*β:V as expected based on chromosome number. Rare I*α*β-LTR activity on V-chromosomes at ∼3 Mya is consistent with homeologous exchange between V and I*α*β. **f)** Scenario for the evolution of octoploid strawberry from diploid progenitors, showing progressive addition of subgenomes and resulting intervals of shared transposon activity.

The clear association of one of the octoploid subgenomes (“V”) with the diploid woodland strawberry *F. vesca* is well-established based on various markers.{Tennessen 2014; Sargent 2016; Edger 2019} At first glance it may appear from the clustering atop **Fig. 1a** that *F. iinumae* is equally related to the other three subgenomes. It is important to realize, however, that the hierarchical clustering of **Fig. 1a** is not a phylogram, but rather an indication of shared repetitive content, which can occur by either (1) shared ancestry, or (2) coexistence in an earlier polyploid progenitor. Since one of the three non-V subgenomes is more closely related to *F. iinumae* based on protein-coding gene phylogeny{Edger 2019, Edger 2020} and whole genome alignment {Liston 2020} we follow standard nomenclature and refer to this as the “I” subgenome. There is no consensus definition of the other two subgenomes{Edger 2019; Spigler 2010; Tennessen 2014; van Dijk et al 2014; Davik et al 2015; Sargent et al 2016} (summarized in {Hardigan, 2020}).

**Fig 1a** shows that these two chromosome sets can be readily separated into ‘α’ and ‘β’ subgenomes based on repetitive content. The subgenomes ‘α’ and ‘β’ are an alternate partitioning of the chromosomes belonging to the ‘nipponica’ and ‘viridis’ subgenomes defined by Edger et al. These two chromosome sets (α plus β, equivalent to ‘nipponica’ plus ‘viridis’) are enriched for 102 13-mer markers relative to I and V. Importantly, however, α chromosomes are differentiated from their homeologous β counterparts by 39 α-enriched 13-mers. We found no 13-mers satisfying the above conditions that are enriched on the β subgenome, which is therefore defined by its deficit of α- (and I-) enriched sequences. **Fig. 1b** shows the chromosomal distribution of V-, I-, α-, αβ- and Iαβ-enriched 13-mers for the first homeologous quartet of octoploid strawberry; the other six quartets are shown in **Ext. Data Fig. 2**. While each chromosome can be assigned to a subgenome based on its overall 13-mer content, the occasional occurrence of markers from one subgenome in localized patches on other chromosomes is consistent with low levels of homeologous exchange, consistent with evidence from protein-coding genes and whole-genome alignment.{Edger 2019; Edger 2020; Liston 2020}

Repetitive content also allows us to evaluate alternative subgenome hypotheses in an unbiased manner by analysis of its variance within vs among subgenomes (ANOVA;**Suppl. Note 2**). **Figs. 1c,d** show “volcano plots” of the repetitive 13-mers content between hypothesized subgenomes, similar to methods used for differential gene expression.{Tukey 1949; Cui 2003} **Fig. 1c** shows that 92 13-mers support our α-β subgenome hypothesis with a Tukey-range-test p-value less than 10^−7^ (p < 0.05 with Bonferroni correction). These include the 39 13-mers of **Fig. 1a** that have two-fold enrichments in all quartets. In contrast, the ‘nipponica’-’viridis’ subgenome hypothesis{Edger 2019} is not supported by any repetitive 13-mers (**Fig. 1d**). This analysis also strongly supports other pairwise differences (**Ext. Data Fig. 3**).

The grouping of subgenomes I, α, and β in **Figures 1a,b**, and **Ext. Data Figs. 1 and 2** arises from recent shared transposon activity. We inferred the timing of this activity from the sequence divergence between 5’- and 3’ long terminal repeats (LTRs){Dangel 1995; Ma 2004}. The divergence of the crown group *Fragaria* has been estimated to be 8 Mya{Qiao 2016} or 3.9 Mya {Dillenberger 2018} (**Supplementary Note 3**). The extensive shared Iαβ activity peaks at divergence corresponding to ∼3 Mya (**Fig. 1e**). This in turn suggests that these three subgenomes were united in a hexaploid prior to the merger with the V subgenome. Combining these data with phylogeny from whole genome alignment{Liston 2020} yields the scenario for the evolution of octoploid strawberry from V, I, α, and β progenitors shown schematically in **Fig. 1f**. We note that the accelerated rate of sequence change in octoploid relative to diploid strawberries, coupled with possible extinction of relevant dipolods, suggest possible explanations for the failure of protein-coding phylogeny to identify the α and β subgenomes (**Suppl. Note 4**).

Edger et al. find that the V subgenome is “dominant,” having lost fewer genes and maintaining on average more robust gene expression than the other subgenomes. This “subgenome dominance” is hypothesized to arise from the resolution of genetic and epigenetic conflicts.{Garsmeur 2014; Edger 2019} Based on **Fig. 1f**, however, we suggest an alternative explanation arising from the neutral expectation that redundant genes are often lost and/or evolve disrupted expression.{Lynch 2007} According to **Fig. 1f**, the Iαβ subgenomes had already evolved under millions of years of redundancy in the hexaploid progenitor prior to the addition of V; conversely, the V subgenome was intact when it was combined with the hexaploid. It follows that the V subgenome of octoploid strawberry is expected to have suffered less disruption (due to gene loss or altered expression) than its Iαβ counterparts simply based on timing. This neutral effect of redundancy and the subgenome conflict hypothesis are not mutually exclusive, but may be difficult to distinguish.

## Extended Data Figure Legends

**Extended Data Figure 1.**
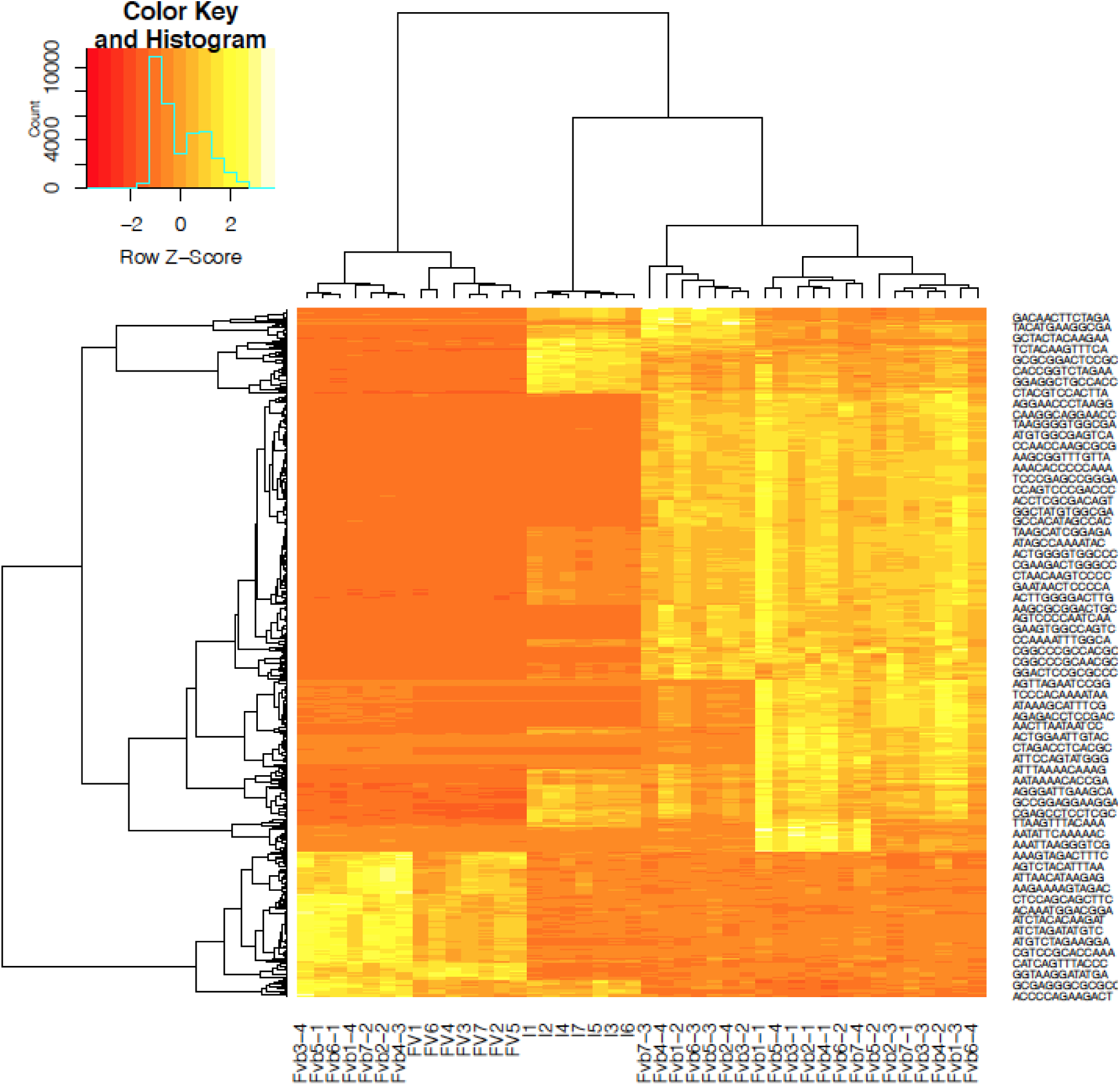
Partition of octoploid strawberry into subgenomes. Hierarchical clustering of chromosomes based on the complete set of 829 repetitive chromosomes described in the text. (Fig 1a shows a subset.)

**Extended Data Figure 2.**
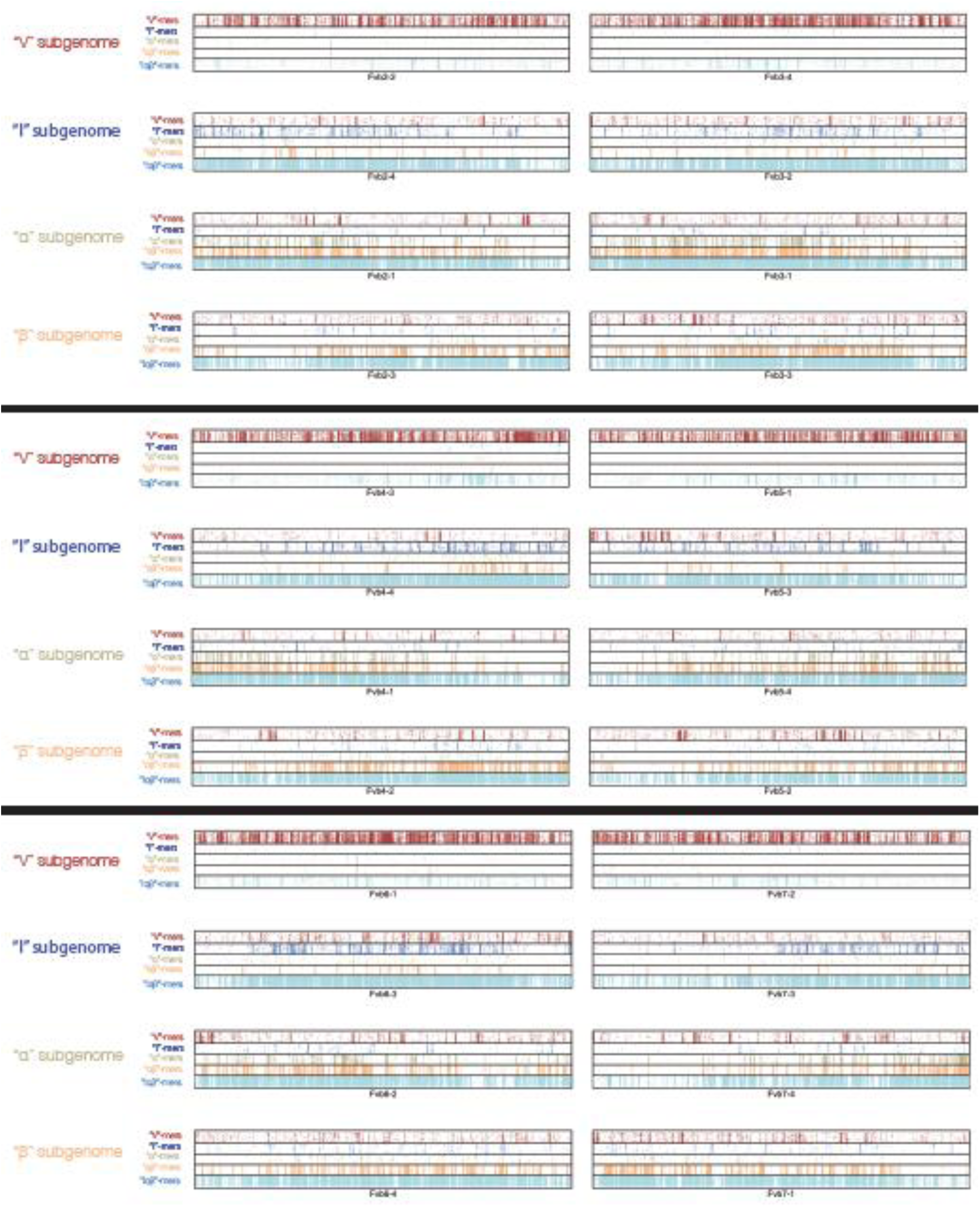
Chromosomal distribution of subgenome enriched 13mers. Distribution of four classes of 13-mer (V, I, *α, α*β, I*α*β) for homeologous quartets 2-7 of octoploid strawberry. Short regions of homeologous exchange are evident.

**Extended Data Figure 3.**
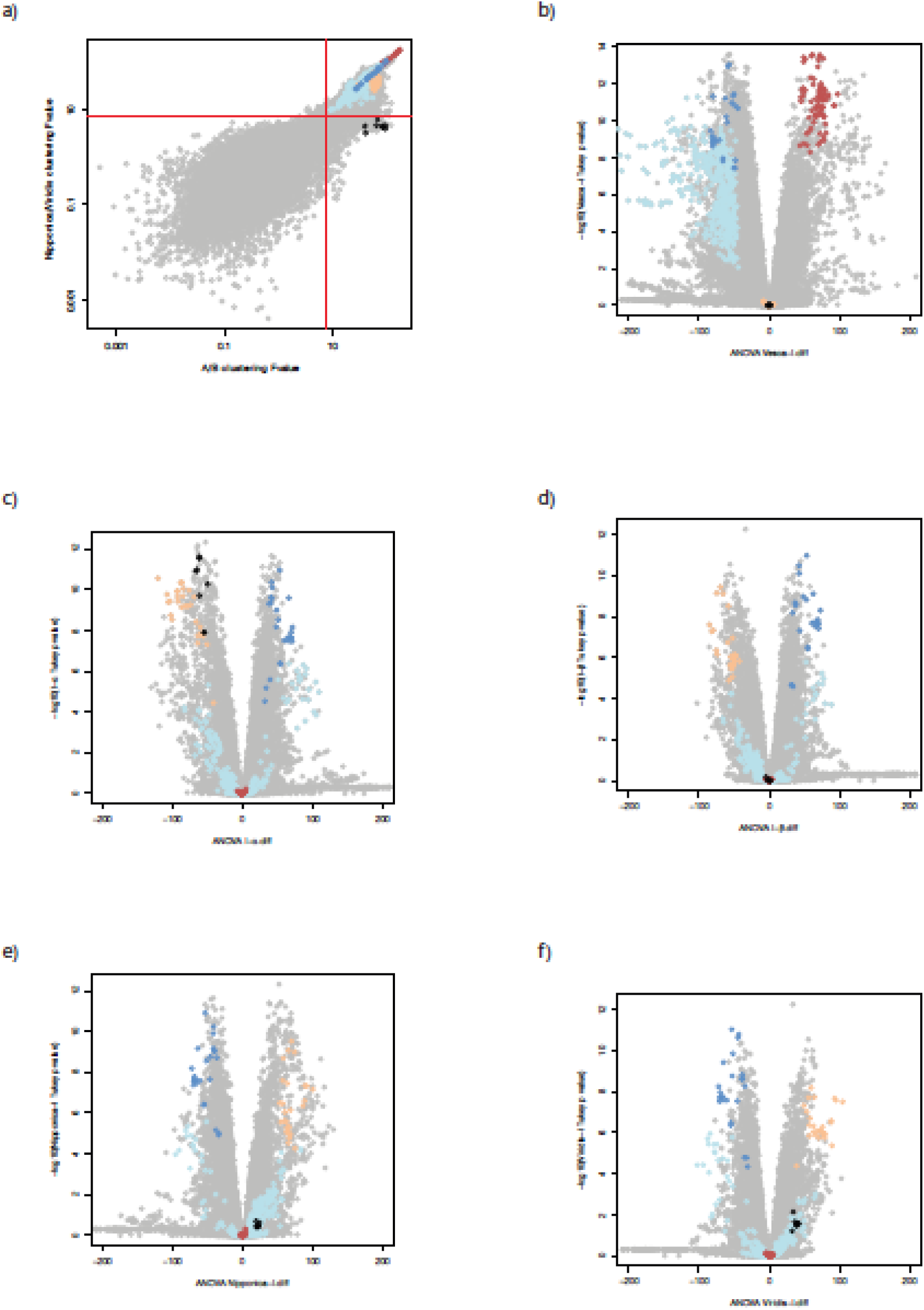
(a) Comparison of ANOVA F value between the *α*/β clustering (x-axis) and the nipponica-viridis clustering (y-axis). Red lines are drawn at F=7.55446, the significant value for *α*=0.001. Black dots (*α*-13mers) are part of a larger cloud that pass our significance threshold, but fail theres. (b-f)Volcano plots demonstrating significant differences between (b) V and I subgenomes, (c) I and *α*, (d) I and β, (e) I and ‘nipponica’, (f) I and ‘viridis’, where ‘viridis’ and ‘nipponica’ are defined in Edger et al. 2019. (Contrasts between V and *α*/β and ‘viridis’/’nipponica’ not shown; these are comparable to the V/I contrast, since V is highly differentiated relative to I, *α*, and β as seen in **Figure 1a**).

**Ext. Data Fig. 4.**
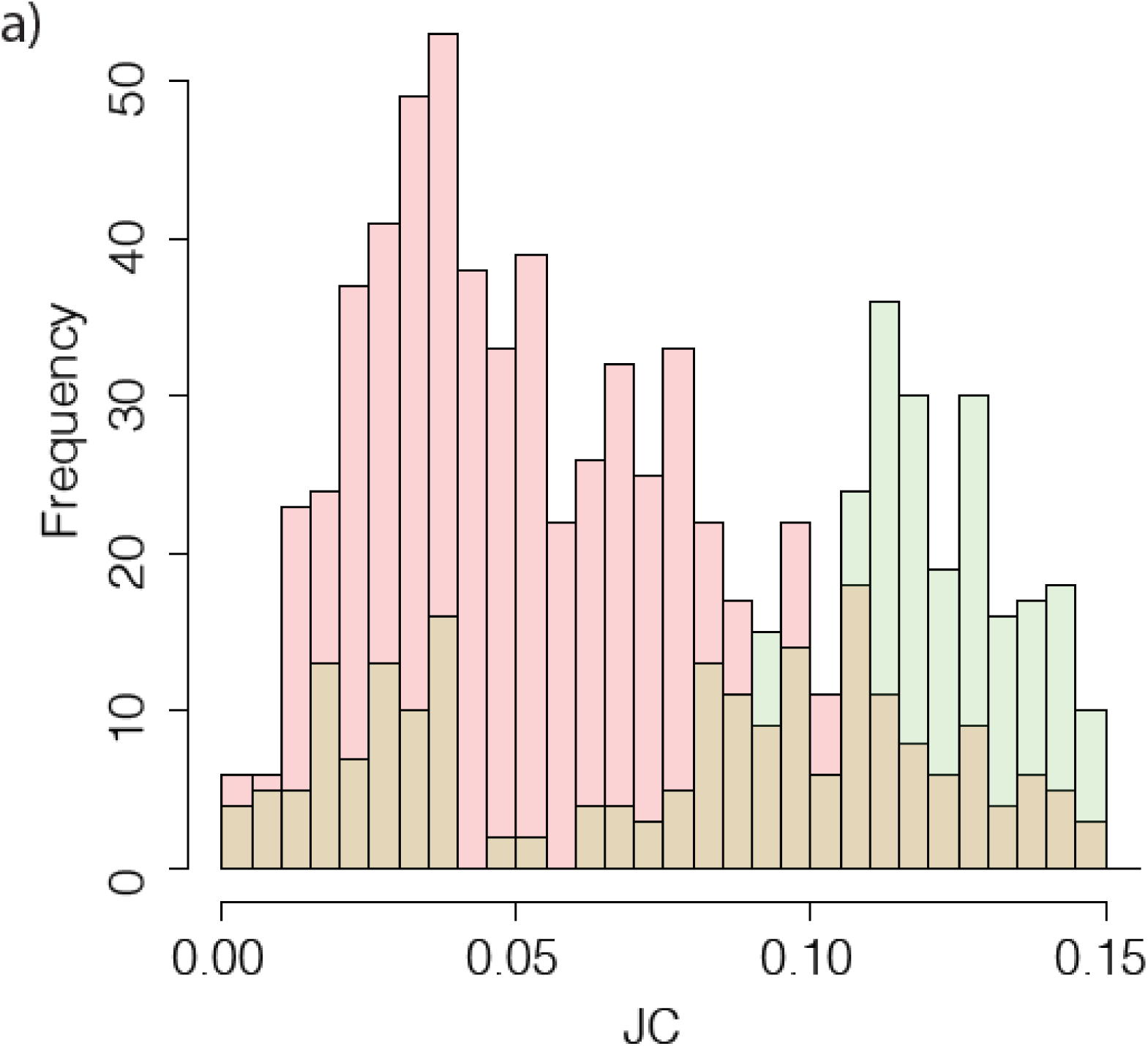
Calibration of sequence divergence vs. time. Histograms of Jukes-Cantor (JC) distance between LTRs of diploid *F. vesca* and octoploid *F. x ananassa*, separated by subgenome type. Mutual best hits from *F. vesca* to I*α*β subgenomes (green) peak at ∼0.11, which we calibrate to 8 My, i.e., the base of the *Fragaria* radiation. Mutual best hits from *F. vesca* to the V subgenome peak more recently ∼0.035, consistent with close relationship between *F. vesca* and this subgenome. Note that there also a small peak in the green curve also ∼0.035, representing likely homeologous exchange (that is, these elements were born on the V subgenome but ended up on I*α*β chromosomes).

## Supplementary Information

**Note 1. Identification of subgenome-specific 13-mers**.

**Note 2. Analysis of variance for subgenome partitions**.

**Note 3. Timing of retrotransposon activity and polypoidy**

**Supplementary Data File 1: Subgenome-specific 13-mers**

829 subgenome-diagnostic 13-mers, with count on each chromosome of *F*. x *ananassa*.

### Note 1. Identification of subgenome-specific 13-mers

We used Jellyfish{Marcais 2011} to count the 13-mers of the octoploid *Fragaria* x *ananassa* genome sequence from Edger *et al*.{Edger 2019} The octoploid strawberry genome (2n=8x=56) is divided into x=7 sets of four homeologous quartets, with each quartet corresponding to a single chromosome of the diploid woodland strawberry *F. vesca*{Edger 2018} and the Japanese diploid *F. iinumae* {Edger 2020}.

We computed the 13-mers that were (1) found in at least 100 copies in the entire octoploid genome, and (2) at least two-fold enriched in one of the four members of each quartet relative to the other members, without regard to the subgenome assignment of Edger *et al*.{Edger 2019} To be considered further, a 13-mer had to be enriched in this manner in all seven quartets. The diploid genomes were not involved in identifying these 13-mers. This computation defined 829 13-mers with potential subgenome contrasts. These 13-mers consistently grouped chromosomes into subgenomes. In the notation of the main text and below, we found 488 13-mers enriched in I, α, and β (relative to V); 175 enriched in V (relative to I, α, and β); 102 enriched in α and β (relative to I and V); 39 enriched in α; and 25 enriched in I (Supplementary Data File 1).

**Figure 1a** shows a hierarchical clustering of the 56 chromosome of *F*. x *ananassa* and the 7 chromosomes of each of the diploids *F. vesca* and *F. iinumae*, with distance function given by 1-Pearson’s correlation of the 13-mer densities (normalized by chromosome length), using the “hclust” function in R. Chromosomes of the diploid *F. vesca* and *F. iinumae* were only available in masked form. Chromosome clustering was insensitive to details (e.g., whether or not the diploids were included), and the same groupings were also reproduced using other clustering methods. At the same time, the 13-mers themselves were also clustered using R, defining (in the notation of the main text and below) 13-mers enriched in the subgenome combinations V, Iαβ, αβ, I, and α, as shown in **Figure 1a**.

Our V and I subgenome assignments agree with previous assignments by Tennessen et al.{Tennessen 2014}, Edger et al.{Edger 2020}, and others (summarized in {Hardigan 2020}):

- **V-subgenome**. We identified the subgenome that groups with diploid *F. vesca* in **Figure 1a** as the “V” subgenome, consistent with Edger et al.{Edger 2019} and previous work based on protein-coding gene phylogeny and similarity (summarized in {Hardigan 2020}). The V subgenome is enriched for V-specific 13-mers includes chromosomes Fvb1-4, Fvb2-2, Fvb3-4, Fvb4-3, Fvb5-1, Fvb6-1, and Fvb7-2.
- **I-subgenome**. Of the three subgenomes that group with *F. iinumae* in **Figure 1a**, one of them is consistently described as the “I” subgenome by multiple authors, based on protein-coding gene phylogeny and similarity{Tennessen 2014; Edger 2019} (see summary in {Hardigan 2020}). The I subgenome is enriched for I-specific 13-mers, and also for 13-mers enriched in Iαβ. The I subgenome includes chromosomes Fvb1-2, Fvb2-4, Fvb3-2, Fvb4-4, Fvb5-3, Fvb6-3, and Fvb7-3.

The remaining two sets of chromosomes were recognized by Tennessen et al.{Tennessen 2014} as (1) closer to each other than to I or V, and (2) closer to I than to V. They have been separated into two sub-genomes by several groups (summarized in {Hardigan 2020} typically based on their similarity to *F. iinumae*, which is recognized as a weak criterion (particularly if these two subgenomes are sister to one another, and so phylogenetically equidistant from *F. iinumae* as suggested by Tennessen et al.{Tennessen 2014}. Edger et al.{Edger 2019; Edger 2020} partitioned these fourteen chromosomes into two sets based on their protein-coding similarity to *F. nipponica* and *F. viridis*, but this was called into question by Liston et al.{Liston 2020}.

We find no repetitive 13-mers whose distribution supports the ‘nipponica’ and ‘viridis’ subgenome assignments, but do find a well-supported an alternate subgenome partition into α and β subgenomes that is distinct from previous studies:

- **The α-subgenome**. The “α” subgenome is enriched for α-specific 13-mers, and also for 13-mers enriched in Iαβ and αβ. It is composed of chromosomes identified by Edger et al. as belonging to the hypothesized ‘nipponica’ or ‘viridis’ subgenomes based on protein-coding similarity and phylogenetic analysis in comparisons with these and other diploids. The α-subgenome includes chromosomes Fvb1-1, Fvb2-1, Fvb3-1, Fvb4-1, Fvb5-4, Fvb6-2, and Fvb7-4.
- **The β-subgenome**. The “β” subgenome is complementary to α. It is marked by 13-mers enriched in Iαβ and αβ but not the α-specific 13-mers. It is composed of chromosomes identified by Edger et al.{Edger 2019; Edger 2020} as belonging to the “nipponica” or “viridis” subgenomes based on protein-coding similarity and phylogenetic analysis in comparisons with these and other diploids. The β subgenome includes chromosomes Fvb1-3, Fvb2-3, Fvb3-3, Fvb4-2, Fvb5-2, Fvb6-4, and Fvb7-1.

**Fig. 1b** (chromosome quartet 1) and **Ext. Data Fig. 2** (chromosome quartets 2-7) show the distribution of V, I, α, αβ, and Iαβ 13-mers across each chromosome. In each quartet the chromosomes are shown in the order V, I, α, and β. There are occasional concentrations of unexpected 13-mers, e.g., the V-enriched sequence at the 3’ end of Fvb1-1 (which is otherwise assigned to the α subgenome). Since the 13-mers are produced by transposon activity, they mark the chromosome identity at the time of transposon insertion. Segments with anomalous 13-mers correspond to homeologous exchanges.

We note that although *F*. x *ananassa* is a hybrid of two octoploids, *F. chiloensis* and *F. virginana*, these two North American species diverged after octoploid formation, and are interfertile, as demonstrated by the conventional disomic meiotic map produced by Hardigan et al. {Hardigan 2020}. Thus we expect their subgenome structure to be the same.

### Note 2. Analysis of variance for subgenome partitions

We assessed the significance of different partitions of the *F*. x *ananassa* genome using analysis of variance (ANOVA), considering the normalized counts per chromosome of the 423,429 13-mers that occur at least 100 times in the octoploid genome. Adopting a significant threshold of 0.05, Bonferroni correction for testing these repetitive 13-mers implies significance threshold of p < 10^−7^. We find that 92 13-mers support our α-β subgenome partition, with 91 found more often on α than β, and 1 found more often on β than α. Similarly 545 13-mers support the partition of I relative to α or β (taking the unique list combining I-α and I-β), and 4,020 13-mers support the partition of the V subgenome from I, α, or β.

In order to test for a significant signal in the 13mer data supporting the *nipponica-viridis* subgenomes of Edger et al., we performed ANOVA, using both our grouping of chromosomes into subgenomes (V, I, α, β) and Edger et al.’s (V, I, ‘nipponica’, ‘viridis’). Since we agree on the V and I subgenomes, we expect V, Iαβ, and αβ 13mers to support both clusterings. The α-specific 13mers, however, contradict the *nipponica-viridis* clustering of Edger et al.{Edger 2019, Edger 2020}.

**Figure 1c, d** shows that the α-specific 13-mers (black circles) are significant in our clustering (as expected based on their definition). All 13-mers that occur at least 100 times in the octoploid genome are shown, with 13-mers identified from our two-fold-enrichment-across-all-quartets criterion shown in color and others in gray.

Evidently there are additional 13-mers with significant α-β contrasts that did not meet the stringent two-fold enrichment criterion imposed in **Note 1**. In contrast, we find no 13-mers that are significant in the ‘*nipponica’-’viridis’* grouping. We also performed a Tukey’s range test (implemented in R with the TukeyHSD function to assign statistical significance to each pairwise subgenome comparison for each 13-mer (**Figure 1c, d; Extended Data Figure 3**). We found significant (p<1e-6) differences between all pairwise subgenome comparisons except ‘nipponica’-’viridis’.

### Note 3. Timing of retrotransposon activity and polyploidy

In order to infer the timing of subgenome-associated transposon activity, we identified LTR retrotransposons in the *F*. x *ananassa* genome using LTRHarvest.{Ellinghaus 2008} We found subgenome-specific LTR retrotransposons by overlapping these sequences with the subgenome-enriched 13-mers defined in **Note 1**, and defined families of subgenome-specific LTRs by sequence-based clustering (using alignments with at least 90% length of the longer sequence and 1e-2 e-value cutoff) using all-vs-all BLASTN{Camacho 2008} with all other parameters set to their default values. Since the 5’ and 3’ long terminal repeats (LTRs) are identical at the time of insertion, the sequence divergence of intact 5’/3’ pairs is proportional to the time since insertion.{Dangel 1995; SanMiguel 1998; Ma 2004} We measured 5’-3’ sequence divergence by Jukes-Cantor distance using the ape package in R{Paradis 2018}.

In order to calibrate 5’-3’ LTR sequence divergence to geological time, we reasoned that best hits of LTRs from the diploid *F. vesca* to LTRs of the I*α*β subgenomes of octoploid *F. x ananassa* would represent divergence of ancient LTRs found in the last common ancestor of these genomes, circa 8 Mya at the base of the *Fragaria* radiation.{Qiao 2016} As shown in **Ext. Data Fig. 4**, this distribution peaks at ∼0.11, so we assign the rate of nucleotide substitution in strawberry LTRs to be = 0.11/(2*8 My) = 0.7 × 10^−8^ subs/year, or 1.4 × 10^−8^ for the 3.9 My calibration. We note that the latter approximates the canonical value of 1.3 x x 10^−8^ /yr used by Ma and Bennetzen {Ma 2004} and San Miguel et al {SanMiguel 1998} for use with LTRs of grasses and often used more generally for plants including strawberry.{Edger 2018}.This canonical rate is derived from aligned intergenic regions, which according to Ma et al. evolve twice as fast as silent sites in coding regions. Thus our LTR calibration is consistent with rates found in other grasses.

**Fig. 1e** shows 5’-3’ LTR distances for LTR families with (1) at least 20 members and (2) overlap with at least one I*α*β-enriched 13-mer. Based on these 13-mers we infer that the LTR retrotransposons were active when the I*α*β subgenomes were present in the same nucleus. Using our calibration, the peak of I*α*β specific activity at 0.035 substitutions shown in **Fig. 1e** corresponds to ∼3 My. We note that I*α*β-enriched families could also have been active in a last common ancestor of the I*α*β progenitors. We consider this unlikely because (1) based on trees Liston et al.{Liston 2020} it is likely that the divergence of these progenitors was closer to the root of *Fragaria*, and (2) detectable 5’3’ LTR pairs are more common from recent activity, due to the ongoing mutation and loss of non-genic sequences in plants{Ma 2004}.

Interestingly, we also find a recent uptick in activity (<0.01 subs ∼ 1.5 Mya) in these families that is present in both I*α*β and V subgenomes, in roughly 3:1 proportion. We interpret this activity as arising from reactivation of I*α*β transposons that were silenced in the hexaploid but released from silencing upon octoploid formation. The timing of this activity then roughly corresponds to the formation of octoploid, consistent with other timing estimates for this event based on protein-coding genes.{Njuguna 2013; Edger 2019} Finally, we also see a small peak in 5’3’ distance for I*α*β-type transposons on the V subgenome, roughly coincident with peak activity on I*α*β. We interpret these LTR pairs as having originally been inserted on I*α*β chromosomes during the hexaploid, but having ended up on the V subgenome due to subsequent homeologous exchange in the octoploid. Note that the timing of homeologous exchange does not affect the 5’-3’ LTR divergence, but merely transports the pair to another chromosome. A similar effect is evident in the calibration **Ext. Data Fig 4**.

In a final note regarding timing, we note that in the read alignment based phylogenies of Liston et al.{Liston 2020}, the lengths of the “Camarosa” branches derived from the octoploid subgenomes are consistently longer than the lengths of their sister diploid *Fragaria* branches. Roughly the octoploid sequences are evolving ∼60% faster than the diploids. This is to be expected due to the relaxation of purifying constraints in polyploids, due to redundancy.{Lynch 2007} Although we have identified *α* and β subgenomes based on repeat sequences that are different from the subgenomes inferred by Edger et al.{Edger 2019; Edger 2020}, we have not positively identified the closest diploid lineages to *α* and β. We agree with Tennessen, Liston et al.{Tennessen 2013; LIston 2020} that these are related to *F. iinumae* but cannot be assigned more specifically to any extant diploid lineage. How can our findings be reconciled with the phylogenetic analysis presented by Edger et al.{Edger 2020}? It is plausible that the accelerated evolution of “Camarosa” sequenced has in some way contributed to long-branch attraction, which might draw fast-evolving octoploid sequences towards deeper branching diploids. But this is speculation.

## Notes

### Competing Interest Statement

The authors have declared no competing interest.

